# Habitat restoration and the recovery of metacommunities

**DOI:** 10.1101/2023.02.10.527972

**Authors:** Klementyna A. Gawecka, Jordi Bascompte

## Abstract

Ecosystem restoration is becoming a widely recognised solution to the biodiversity crisis. However, there is a gap between restoration science and practice. Specifically, we lack a theoretical framework which would improve our understanding of ecosystems’ recovery and allow us to optimise restoration design. Here, we narrow this gap by developing spatially explicit metacommunity models and studying the recovery dynamics of communities during restoration. We show that community response depends on how damaged the landscape is prior to restoration, with highly fragmented landscapes imposing greater challenges to community recovery. In such cases, we found that the recovery depends on the type of interaction and the structure of the interaction network. Finally, we demonstrate that community recovery can be maximised with careful spatial planning. When recovering communities composed of antagonistic interactions, restoration should target areas adjacent to the most species-rich sites. In the case of mutualistic communities, the same strategy should be adopted in the short-term, whereas in the long-term, restoration should be extended to sites that improve the overall connectivity of the landscape. Our results highlight the importance of considering interactions between species and spatial planning in restoration projects.

## 1 Introduction

Anthropogenic activities are degrading ecosystems around the world at an alarming rate. This degradation, in turn, accelerates the loss of biodiversity, as well as the ecosystem services on which we rely (Millennium Ecosystem Assessment, 2005; IPBES, 2019). In response to this problem, ecosystem restoration has been recognised as a sustainable and efficient solution (Strassburg et al., 2020; United Nations, 2021), and has been receiving more attention in the recent years (Rovere, 2015; Svenning et al., 2016; Bertolini and da Mosto, 2021). With increasing public awareness, growing support from international organisations, and the current United Nations Decade on Ecosystem Restoration (2021-2030), the opportunities for restoration have never been greater.

Despite this, restoration is still based on case-by-case, trial-and-error practices (Ladouceur and Shackelford, 2021), with outcomes being highly varied and often unsuccessful (Suding, 2011; Moreno-Mateos et al., 2017; Brudvig and Catano, 2021). This unpredictability poses challenges for the design and planning of restoration projects. Thus, understanding and predicting the consequences of restoration is of paramount importance for its success. Moreover, there is a substantial gap between science and practice (Cabin et al., 2010; Miller et al., 2017; Clark et al., 2019). Specifically, we currently lack a general theory of ecological recovery (Török and Helm, 2017; Montoya, 2021) which would enable us to develop a conceptual framework for providing rules of thumb on the most efficient restoration design.

Restoration research has been mainly focused on enhancing biodiversity, primarily of plant species (Brudvig, 2011). However, it is the interactions between species, rather than species diversity itself, that mediate ecosystem functioning and its services (Bengtsson, 1998). Despite their importance, ecological interactions have rarely been considered in restoration research (Kaiser-Bunbury et al., 2010; Brudvig, 2011). A powerful tool for studying ecological interaction is network analysis where nodes represent species and links connecting them represent ecological interactions (Bascompte and Jordano, 2014; Raimundo et al., 2018). When applied to restoration, this network approach has already been shown to provide insights which would have been overlooked if the focus were only on the species (Devoto et al., 2012; Pocock et al., 2012; Mittelman et al., 2021; Gaiarsa and Bascompte, 2022; O’Connell et al., 2022).

Although experimental and observational studies on restoration of ecological interactions can provide invaluable information (e.g., Forup et al., 2008; Ribeiro da Silva et al., 2015; Kaiser-Bunbury et al., 2017), they are restricted to specific conditions, small spatial and short temporal scales (Solé and Bascompte, 2006). Conversely, theoretical approaches allow the study of the effects of restoration at a variety of scales in a time- and resource-efficient manner. Furthermore, they enable complex processes to be disentangled, and thus can guide restoration design (Montoya et al., 2012; Maschinski and Quintana-Ascencio, 2016). Metapopulation theory, in particular, has been invaluable to our understanding of species’ dynamics in space, and has been applied to numerous conservation problems (e.g., Levins, 1969; Lande, 1987; Bascompte and Solé, 1996; Huang et al., 2020). However, previous theoretical studies exploring the consequences of restoration focus on the response of a single species or simple communities (Tilman et al., 1997; Huxel and Hastings, 1999; Meyer, 2019; Gawecka and Bascompte, 2021). Thus, we still lack an understanding of the recovery of entire communities of interacting species (Moreno-Mateos et al., 2020; Bullock et al., 2022; Windsor et al., 2023).

Here, we aim to narrow the science-practice gap by proposing a theoretical framework which employs metacommunity theory and empirical interaction networks, and enables us to study the response of communities to habitat restoration. By developing spatially explicit metacommunity models, we explore the recovery dynamics of mutualistic and antagonistic interaction networks. First, we investigate how the amount of previously destroyed habitat and the structure of interaction networks affect the recovery of local interaction richness (number of interactions in a habitat patch), network similarity (similarity of interactions between local networks) and regional interaction abundance (fraction of patches where an interaction is present). Second, by simulating alternative restoration strategies, we investigate what spatial arrangements of restored sites can enhance the recovery, and provide recommendations on the spatial planning of restoration projects.

## 2 Methods

### 2.1 Ecological networks

We considered communities where species belonging to two different guilds interact either mutualistically or antagonistically, and thus they can be represented as bipartite networks. Examples of such mutualistic networks include plants and their insect pollinators or seed dispersers, whereas antagonistic networks include plants and their herbivores or hosts and their parasites. In both cases, we refer to the lower guild as the resource, and the upper guild as the consumer. In our simulations, we used empirical networks (20 mutualistic and 20 antagonistic, Table S1) from the Web of Life repository (www.web-of-life.es; Fortuna et al., 2014) as the metacommunities or metanetworks (the networks of all species and interactions across the entire pristine landscape, Figure 1A).

**Figure 1:**
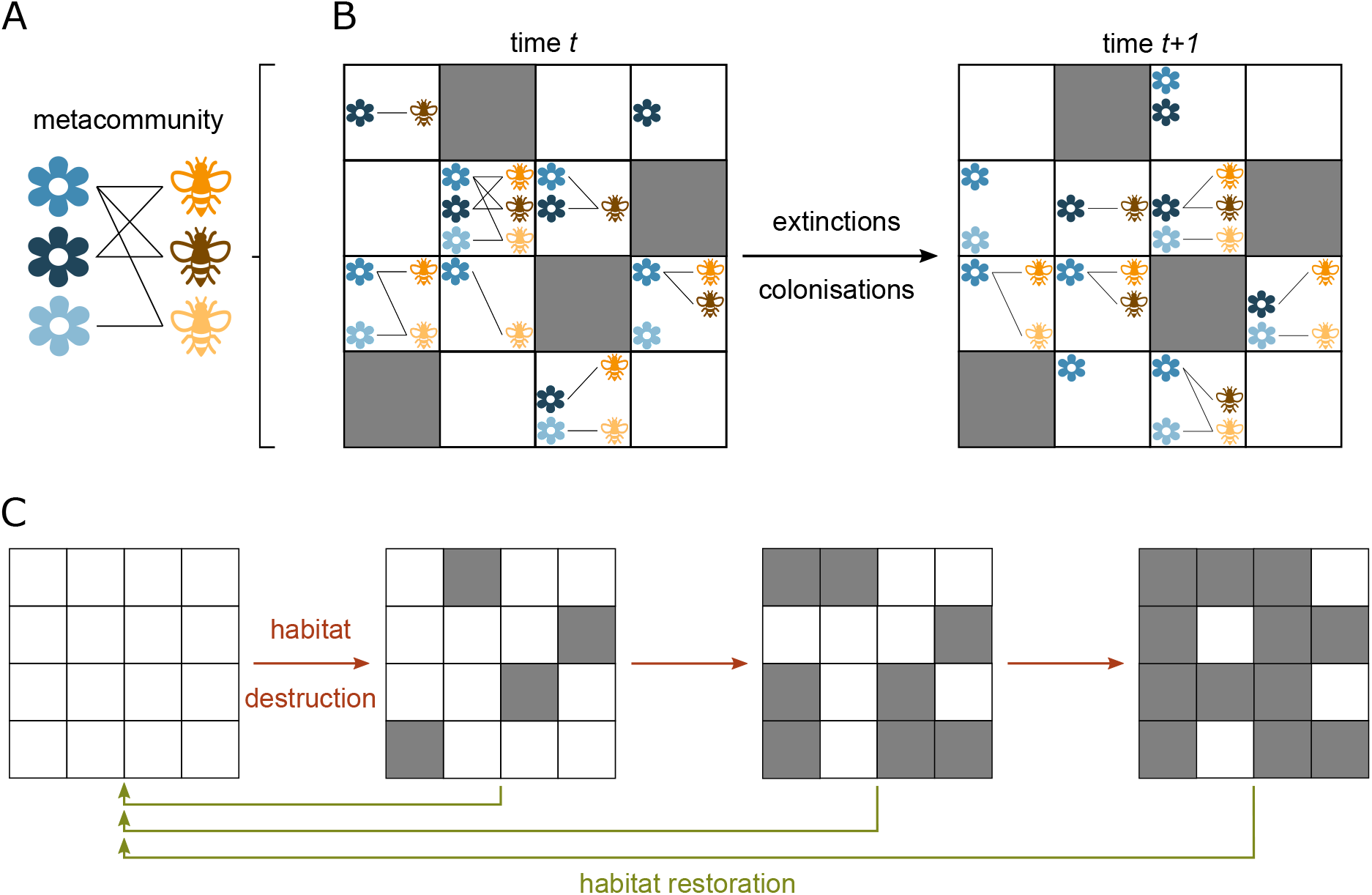
Our spatially-explicit metacommunity model. (A) Metacommunity represented as a metanetwork — network of all species and interactions across the landscape. We use empirical mutualistic and antagonistic bipartite networks as the metanetworks. (B) Landscape containing pristine (white squares) and destroyed (grey squares) patches. Destroyed patches cannot harbour any species, whereas pristine patches can be occupied by the entire metacommunity, or a subset. Metacommunity dynamics are governed by local extinctions and colonisations of neighbouring patches. (C) Simulating habitat destruction followed by its restoration. Each restoration simulation starts at a different point of the destruction process and continues until all patches are pristine again. After destroying or restoring a fraction of patches, we simulate metacommunity dynamics (panel B) until reaching the steady state.

### 2.2 Spatially-explicit metacommunity model

We discretised our landscape into a regular grid of identical square patches, with dimensions of 100 × 100 patches. To simulate the metacommunity dynamics in space, we adopted the patch dynamics perspective (Leibold et al., 2004), whereby patches may be occupied or unoccupied by a species. We assumed that pristine patches can harbour the entire empirical network (i.e., the metanetwork). However, as a result of the stochastic spatio-temporal dynamics, a patch may be occupied by only a subset of the species in the metanetwork, or it may be empty (Figure 1B).

The spatial dynamics were governed by local extinctions and colonisations (Table 1). A species occupying a patch at a given time step had a probability of becoming locally extinct at the next time step. In mutualistic communities, the extinction probabilities of all species were independent of the species composition in the patch (Fortuna et al., 2013). Conversely, the extinction probability of resources in antagonistic communities increased with the number of its consumers present in the patch. As for the antagonistic consumers, the more resources they had in a patch, the less likely they were to become locally extinct.

**Table 1:** Effective extinction, *p*_*e,i*_, and colonisation, *p*_*c,i*_, probabilities of species *i* in mutualism and antagonism.

|  | mutualism | antagonism |
| --- | --- | --- |
| **extinctions** |  |  |
| resource | $p_{e,i} = e_{r,i}$ | $p_{e,i} = 1 - (1 - e_{r,i}) \prod_{j=1}^J (1 - e_{r,i}/j)$ |
| consumer | $p_{e,i} = e_{c,i}$ | $p_{e,i} = e_{c,i} \prod_{j=1}^J (1 - e_{c,i}/j)$ |
| **colonisations** |  |  |
| resource | $p_{c,i} = 1 - \prod_{n=1}^N \prod_{j=1}^J (1 - c_{r,i}/j)$ | $p_{c,i} = 1 - (1 - c_{r,i})^N$ |
| consumer | $p_{c,i} = 1 - \prod_{n=1}^N \prod_{j=1}^J (1 - c_{c,i}/j)$ | $p_{c,i} = 1 - (1 - c_{c,i})^N$ |
| $e_{r,i}$ - intrinsic extinction probability of resource $i$ | | |
| $e_{c,i}$ - intrinsic extinction probability of consumer $i$ | | |
| $c_{r,i}$ - intrinsic colonisation probability of resource $i$ | | |
| $c_{c,i}$ - intrinsic colonisation probability of consumer $i$ | | |
| $J$ - number of interaction partners $j$ of species $i$ | | |
| $N$ - number of neighbouring patches which are suitable sources for colonisation | | |

If a species were absent from a patch in a given time step, it had a chance of colonising it from one of the eight nearest neighbours (i.e., assuming Moore’s neighbourhood) in the next time step. The colonisation probability of the resource in mutualism increased with the number of consumers with which they shared a patch, and the number of such patches. This assumed consumer-mediated dispersal (Fortuna et al., 2013). The colonisation probability of the mutualistic consumers increased with the number of neighbouring patches where they were present and the number of their resources in the patch to be colonised (Fortuna et al., 2013). In antagonistic communities, the colonisation probability of both resources and consumers increased with the number of neighbouring patches where they were present, but was independent of the species composition.

Here, we present the results of simulations with one set of intrinsic extinction and colonisation probabilities (*e*_*r,i*_ = *e*_*c,i*_ = 0.2, *c*_*r,i*_ = *c*_*c,i*_ = 0.1 for mutualism, and *e*_*r,i*_ = 0.1, *e*_*c,i*_ = 0.4, *c*_*r,i*_ = *c*_*c,i*_ = 0.1 for antagonism, see Table 1). These values ensure regional coexistence of all species in a pristine landscape, although not necessarily in each patch. Note that while varying these parameters changes the results quantitatively, it does not affect the qualitative outcomes (Figures S4-S6).

### 2.3 Simulations

The first simulation stage was destruction of habitat (Figure 1C). Starting with a landscape where all patches were pristine and all harboured the entire metacommunity, we simulated the extinction-colonisation dynamics until we reached a steady state where the regional abundance of each species (the number of patches they occupy) no longer changed between time steps. We then destroyed the landscape by randomly selecting 500 pristine patches (5% of the landscape) and changing their state to ‘destroyed’. We assumed that all species in those patches immediately became locally extinct, and that these patches could not be recolonised. This was followed by simulating metacommunity dynamics over time until a new steady state was reached, and the process continued until the entire landscape was destroyed. This resulted in steady state landscapes with habitat loss varying from 0% (fully pristine) to 100% (fully destroyed) in steps of 5%.

The second stage involved simulating habitat restoration (Figure 1C). For each steady state landscape from the destruction stage (21 landscapes varying from 0 to 100% habitat loss), we restored it in steps of 5% by selecting 500 destroyed patches and changing their state to ‘pristine’. Thus, we assumed that restoration of the habitat in those patches was immediate. At each step, we simulated the metacommunity dynamics until reaching a new steady state. Starting restoration at different fractions of habitat loss allowed us to assess the effect of the amount of previously destroyed habitat on the restoration outcome.

Furthermore, we investigated the effect of different spatial restoration strategies on community recovery. In the ‘random’ strategy, the patches to be restored were always selected at random, as in the destruction stage. In the ‘nonrandom’ strategy, we prioritised restoration of patches adjacent to pristine patches harbouring the largest number of species. This resulted in the landscape being restored in clusters centred around the most species-rich areas. We also explored a third strategy, ‘alternative nonrandom’ where we selected destroyed patches which were adjacent to pristine patches occupied by at least one species. However, as the results of the two ‘nonrandom’ strategies are qualitatively similar, we present the latter in Supporting Information (Figures S7-S10).

We repeated these simulations with each of the 40 empirical networks as the metanetworks. All simulations were performed with Julia version 1.4.2 (Bezanson et al., 2017), whereas data visualisation was done with R version 3.6.2 (R Core Team, 2020).

### 2.4 Data analysis

To asses the effects of habitat destruction and restoration, we analysed three measures covering different spatial scales: local interaction richness, network similarity, and regional interaction abundance. We defined the local interaction richness as the fraction of all interactions in the metanetwork present in a given patch. To quantify the similarity of the local networks, we measured *β*-diversity. At each fraction of habitat loss during destruction and restoration, we sampled 100 patches. Then, using the Whittaker measure (Whittaker, 1960) in the *betalink* package in R (Poisot et al., 2012), we computed the interaction similarity (1 − *β*) between all pairwise combinations of the sampled patches (hereafter ‘network similarity’). For local interaction richness and network similarity, we obtained their mean values across all patches in the landscape where a network was present. Lastly, for each interaction in the metanetwork, we calculated its regional abundance defined as the fraction of all patches in the landscape where it was present.

To quantify the response of communities to restoration in different scenarios, we adopted a measure of ‘restoration efficiency’ (*R*) proposed by Gawecka and Bascompte (2021). Although it was developed for species abundance, here, we apply it to the three measures described above (local interaction richness, network similarity and regional interaction abundance). Restoration efficiency is defined as:

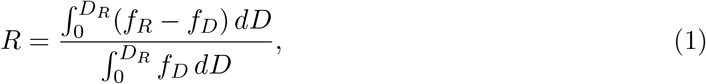

where *D*_*R*_ is the fraction of habitat loss at which restoration starts, whereas *f*_*D*_ and *f*_*R*_ are the curves of the measure under consideration (local interaction richness, network similarity or regional interaction abundance), as a function of habitat loss, *D*, during habitat destruction and restoration, respectively. Thus, *R* represents the difference between the areas enclosed by the two curves and the habitat loss axis between 0 and *D*_*R*_, and is normalised by the area below the habitat destruction curve. Note that *R* summarises the outcome of the entire restoration process (i.e., from the fraction of habitat loss at which restoration begins, until the entire landscape is pristine). Negative values of *R* indicate that, overall, the measure under consideration is lower during restoration than during destruction. We refer to this type of response as ‘restoration lag’ (Tilman et al., 1997). Conversely, positive values of *R* are referred to as ‘restoration boost’ (Gawecka and Bascompte, 2021). *R* = 0 indicates that the overall response during restoration is the same as during destruction. Finally, *R* = −1 corresponds to a scenario where the community, despite restoration, became globally extinct.

We analysed the structure of the empirical networks (i.e., the metanetworks) by considering connectance, nestedness and modularity. Connectance is the proportion of all possible interactions that are realised in the network. Nestedness is a pattern whereby there is a core group of generalist species interacting with each other, and extreme specialists interacting only with the generalist species. We quantified it using the measure proposed by Fortuna et al. (2019) which is equivalent to the NODF metric (Almeida-Neto et al., 2008) but does not penalize the contribution to nestedness of species with the same number of partners. Modularity indicates to what extent species form modules with many interactions among the species in their modules and very few with species in other modules. We use the *igraph* package in R with a multi-level modularity optimisation algorithm (Blondel et al., 2008) to measure it. As the three descriptors of network structure are highly correlated (Fortuna et al., 2010), we combined them into a single measure using a principal component analysis. The first principal component (PC1) explained 85% of the variance, and thus, we adopted it as the compound structure measure. It was positively correlated with connectance and nestedness (95% and 91%, respectively), and negatively correlated with modularity (−91%).

We then investigated which properties of the metanetwork or the interaction affect the restoration efficiency, *R*. For the network level restoration efficiency measures (local interaction richness and network similarity), we used linear models with metanetwork structure (PC1) and metanetwork size (total number of species in the metanetwork) as the explanatory variables. For restoration efficiency of regional interaction abundance, we used linear mixed models in the

*lme4* package in R. We chose metanetwork structure, metanetwork size, interaction extinction threshold (fraction of habitat loss at which that interaction becomes globally extinct), and interaction degree (sum of degrees of the two species involved in the interaction, Gawecka et al., 2022) as the fixed effects, and metanetwork ID as the random effect. We fitted these models for each restoration simulation (i.e., restoration starting at different fractions of habitat loss), scaling the explanatory variables such that their slope parameter estimates are comparable for a given restoration simulation (Schielzeth, 2010). We then extracted the slope parameter estimate for each explanatory variable. To assess its significance, we chose a conservative p-value of 0.01, as well as computed the *R*^2^.

## 3 Results

### 3.1 Community’s response to habitat restoration

Restoration of the landscape generally results in increasing local interaction richness, local networks becoming more similar to each other, and increasing regional interaction abundance (Figure 2, blue lines). However, the exact response during restoration depends on the fraction of habitat loss at which restoration begins. When restoration starts in low/moderately destroyed landscapes (less than ∼50% habitat loss), community’s response is non-hysteretic, meaning it occurs through the same path as the behaviour during habitat destruction (compare grey and light blue lines in Figure 2). Thus, our restoration efficiency metric is zero (Figure 3).

**Figure 2:**
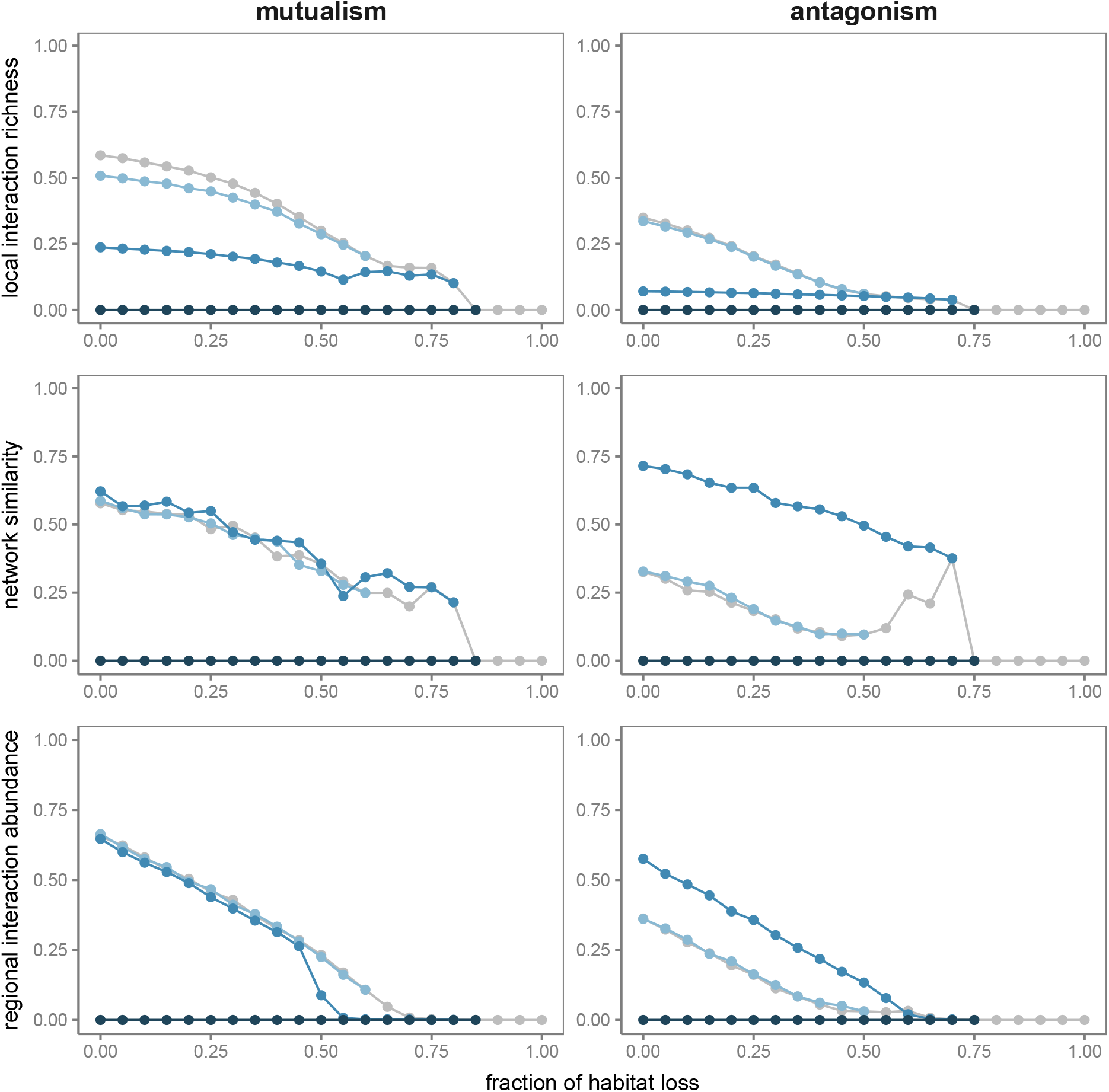
The response to habitat destruction (grey) and ‘random’ habitat restoration (blues). The panels show: mean local interaction richness (top), mean similarity of local networks (middle), and regional abundance of one interaction (bottom). Panels on left and right show the results of simulations with one mutualistic (M_PL_006) and one antagonistic (A_HP_047) network, respectively, as examples. Shades of blue correspond to restoration simulations starting at different fractions of habitat loss (lighter to darker blues: 60, 80 and 85% for the mutualistic network, and 50, 70 and 75% for the antagonistic network).

**Figure 3:**
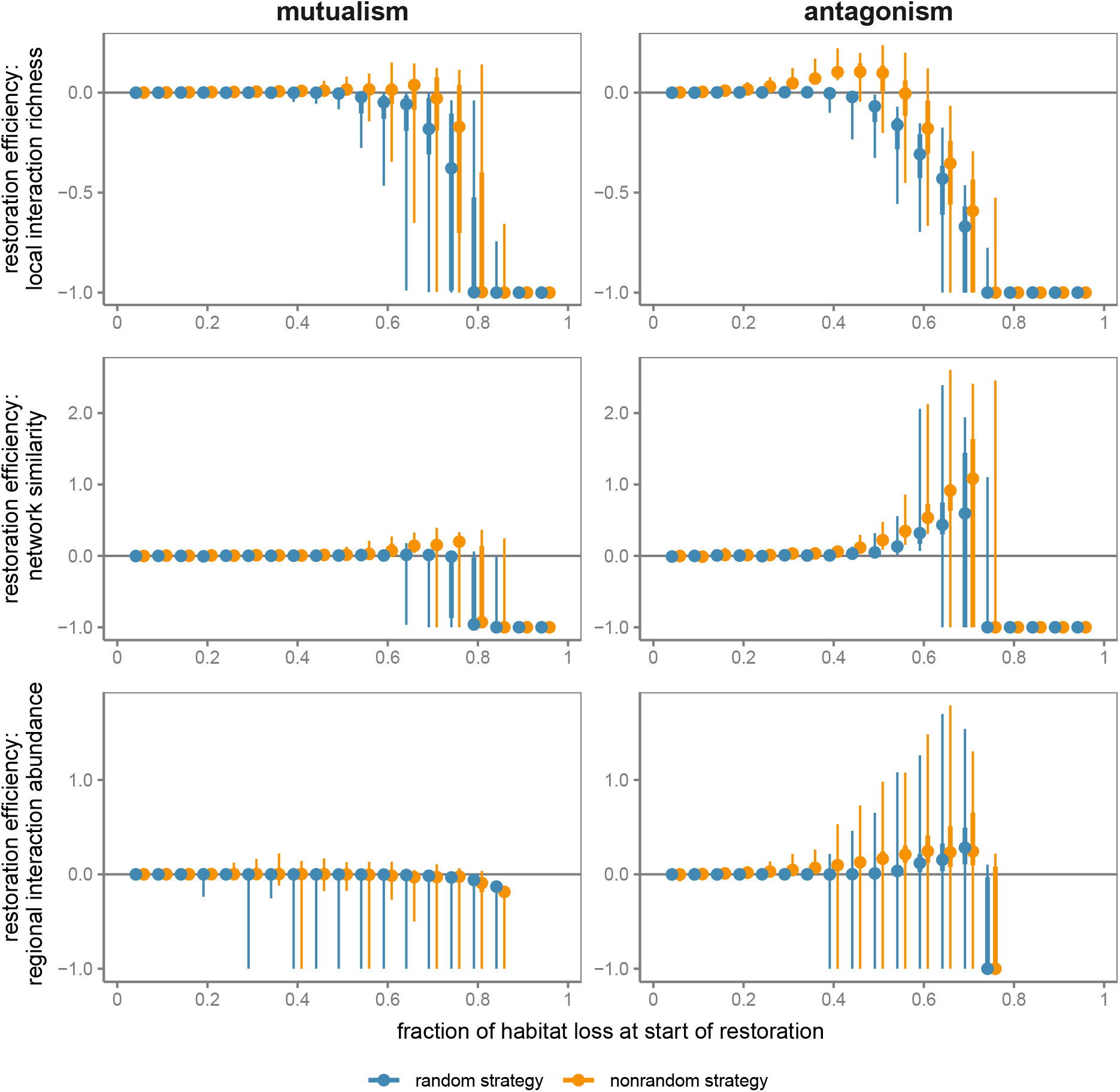
Restoration efficiency as a function of the fraction of habitat loss at which restoration begins. The panels show restoration efficiency of: mean local interaction richness (top), mean similarity of local networks (middle), and regional interaction abundance (bottom). Points, thicker vertical lines and thinner vertical lines represent the median, interquartile range, and minimum and maximum values, respectively, across all mutualistic (left) or antagonistic (right) networks and interactions. ‘random’ and ‘nonrandom’ restoration strategies are shown in blue and orange, respectively. Positive restoration efficiency values indicate a ‘restoration boost’ (i.e., the quantity is, on average, higher during restoration than during destruction), whereas negative restoration efficiency values imply a ‘restoration lag’ (i.e., the quantity is, on average, lower during restoration than during destruction).

As the fraction of habitat loss at which restoration begins increases, there is an increasing restoration lag (negative restoration efficiency) in the recovery of the local interaction richness (Figure 3, top panels). This is largely due to some species, and thus interactions, becoming extinct from the landscape during habitat destruction, such that the local interaction richness cannot recover to its value in a pristine landscape. However, the increase in the local interaction richness is gradual at the initial stages of restoration (intermediate blue lines in Figure 2, top panels). In other words, the local interactions do not reach their potential richness immediately, suggesting that part of the lag is driven by factors other than species extinctions. Moreover, we evaluated the effects of the metanetwork properties on the magnitude of the lag in the local interaction richness recovery (Figure 4, top panels). In the case of mutualism, we found that larger, more connected and nested metanetworks experience a smaller lag than smaller and more modular ones. However, in antagonism, the effects of the metanetwork structure or size are not substantial.

**Figure 4:**
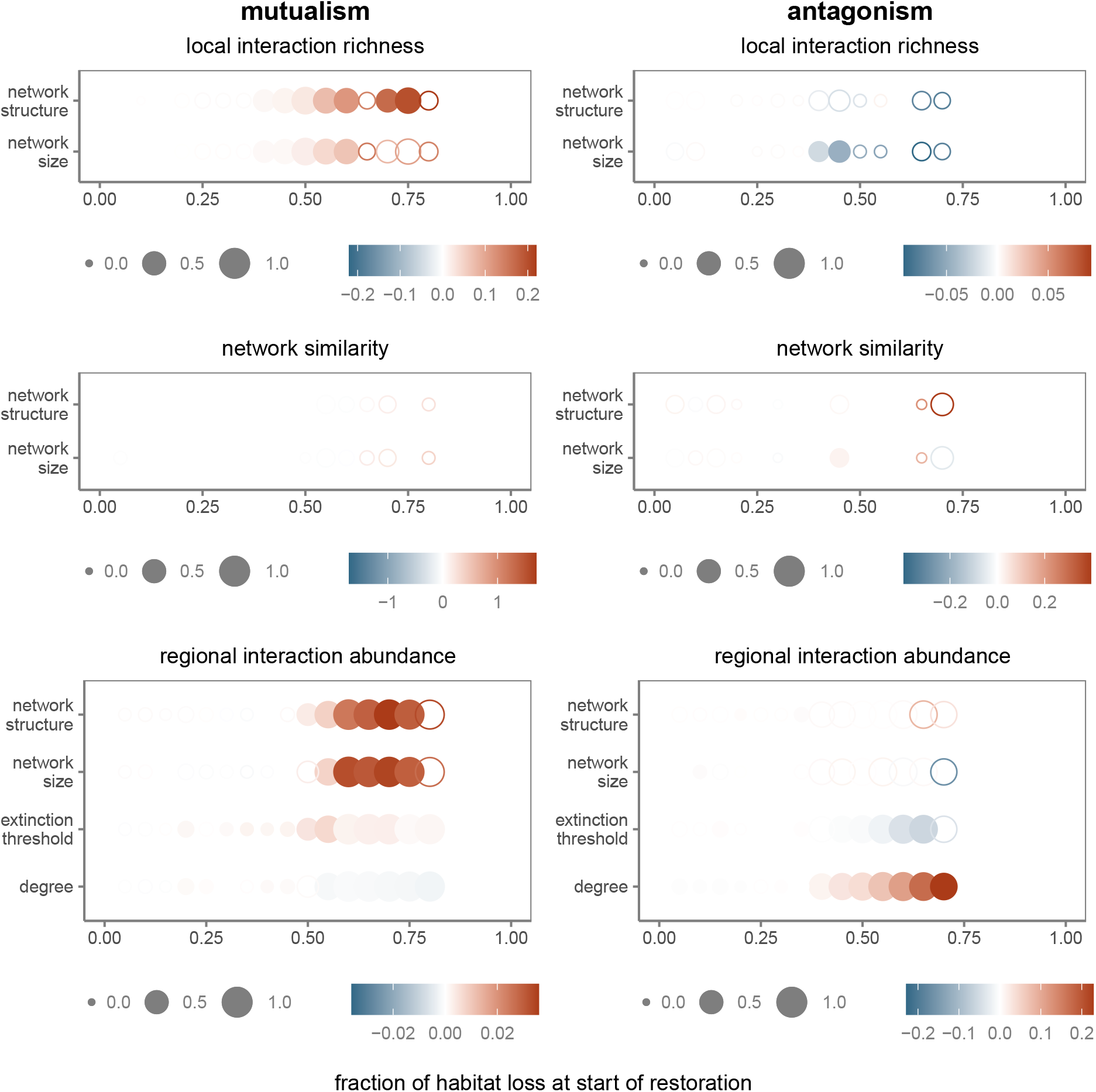
The effect of metanetwork and interaction properties on restoration efficiency of mean local interaction richness (top), mean similarity of local networks (middle), and regional interaction abundance (bottom). For local interaction richness and network similarity, we fitted linear models with metanetwork structure and metanetwork size as the explanatory variables. For regional interaction abundance, we fitted linear mixed models with metanetwork structure, metanetwork size, interaction extinction threshold and interaction degree as the explanatory variables. We fitted the models to the results of simulations with mutualistic (left) and antagonistic (right) networks, and ‘random’ restoration starting at different fractions of habitat loss (see Figures S1 and S8 for ‘nonrandom’ strategies). The size of the points indicates the *R*^2^, full and empty circles correspond to significant (p-value < 0.01) and non-significant effects, respectively, and the colours show the slope parameter estimate. The explanatory variables are scaled, meaning that the magnitudes of their slope parameter estimates are comparable for each interaction type and each restoration simulation.

In terms of the similarity of the local networks, its restoration efficiency reduces abruptly from 0 to -1 in mutualism as the starting fraction of habitat loss increases (Figure 3, middle left panel). In other words, mutualistic network composition across the landscape is as heterogeneous during restoration as during destruction, unless restoration begins in highly fragmented landscapes where the communities cannot recover and become globally extinct (Figure 2, middle left panel). However, antagonistic communities experience a boost in the similarity of local networks when restoration begins in highly fragmented landscapes and the local communities are very small (intermediate blue lines in Figure 2, middle right panel). In these scenarios, restoration results in more homogeneous community composition than in pristine landscapes with the entire metacommunity present. We found no significant effects of metanetwork structure or size on the restoration efficiency of network similarity (Figure 4, middle panels).

At the regional scale, we also found differences between the recovery of mutualistic and antagonistic interactions when restoration starts in highly fragmented landscapes (Figure 3, bottom panels). While the regional abundance of mutualistic interactions exhibits a lag, antagonistic interactions experience a boost. The lag indicates that, overall, the abundance during restoration is lower than during destruction, whereas the opposite is true for the boost (intermediate blue lines in Figure 2, bottom panels). Furthermore, we found that the magnitude of this lag in mutualistic communities is affected by the properties of the metanetwork, with larger, more connected and more nested metacommunities having smaller lags (Figure 4, bottom left panel). Conversely, in antagonistic communities, restoration efficiency is driven by the properties of the interaction, where interactions with higher degree (e.g., interactions between generalist consumers and generalist resources) and lower extinction thresholds have larger boosts (Figure 4, bottom right panel). However, note that the effect of interaction degree is stronger than that of extinction threshold.

### 3.2 Effect of restoration strategy

Next, we investigate the effect of spatial restoration strategies on the community’s response. In the ‘random’ strategy, we restored patches in a random sequence, whereas in the ‘nonrandom’ strategy, we prioritised restoration of patches adjacent to those harbouring the most species-rich communities.

Overall, we found that the ‘nonrandom’ strategy results in a higher restoration efficiency of all three measures (local interaction richness, network similarity and regional interaction abundance), producing either smaller lags or larger boosts, and reducing the likelihood of communities becoming globally extinct (Figure 3). The benefits of the ‘nonrandom’ strategy are particularly evident in antagonism, indicating that restoration actions should always target species-rich areas.

However, considering the regional abundances of mutualistic interactions, the differences between restoration efficiency of the two strategies are not substantial (Figure 3, bottom left panel). Exploring the response of individual interactions, we found that the strategy that produces higher regional abundance depends on the amount of restored habitat (Figure 5; see top panels for all interactions in all networks, and middle and bottom panels for an example response of a single interaction). At the initial stages of restoration (i.e., at high fractions of habitat loss), the ‘nonrandom’ strategy achieves higher regional abundances than the ‘random’ strategy. However, once some habitat has been restored, it is the ‘random’ strategy that tends to result in higher abundances of mutualistic interactions. In other words, it is better to start by restoring patches adjacent to where communities are still present. However, as restoration continues, switching to restoring patches at random may be more beneficial. Note that this applies to scenarios where restoration begins in highly fragmented landscapes. When restoring low/moderately destroyed landscapes, community’s response tends to be non-hysteretic (Figure 3) and there is no difference between the strategies (Figure S2-S3).

**Figure 5:**
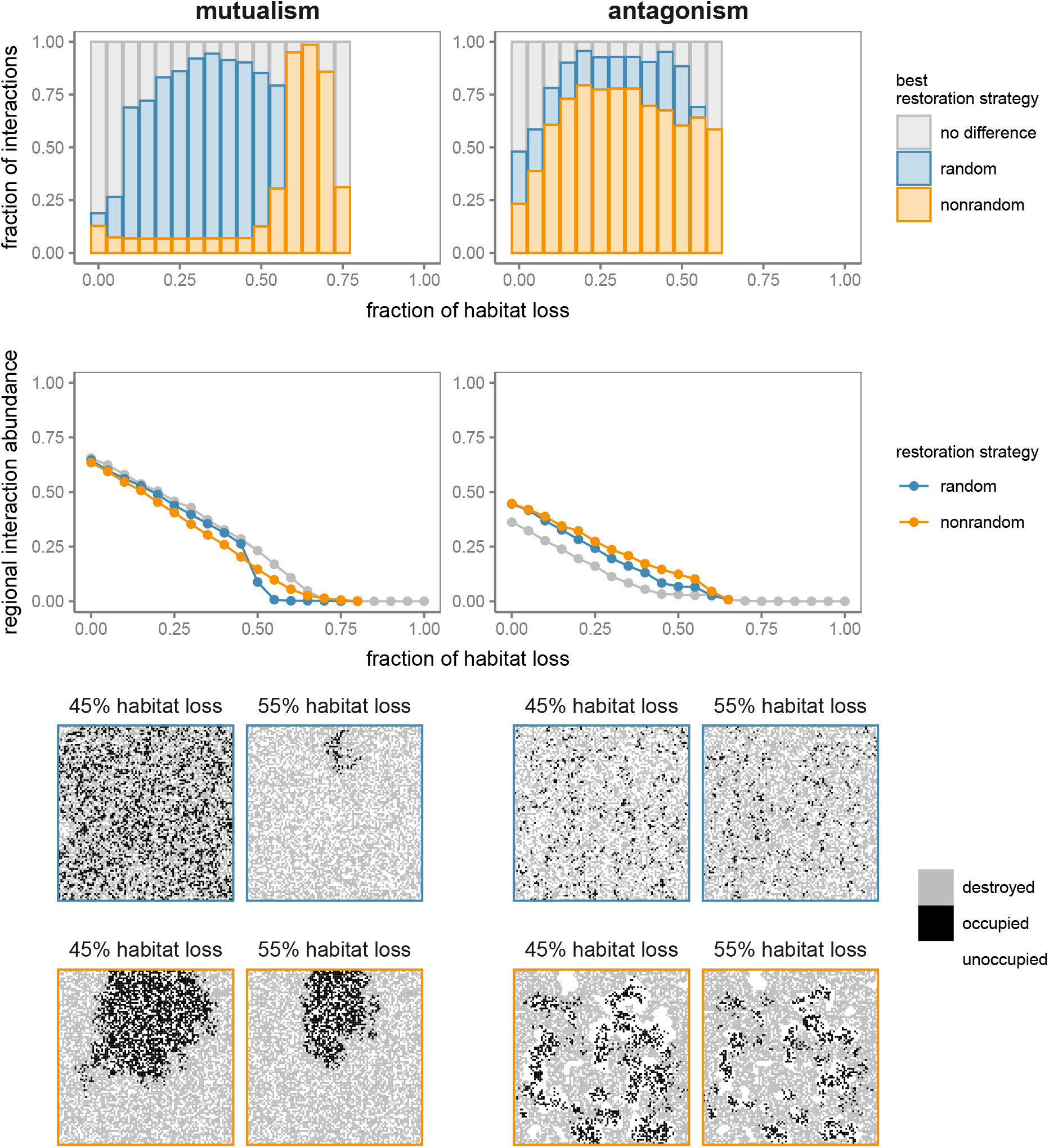
Comparison of ‘random’ (blue) and ‘nonrandom’ (orange) restoration strategies in mutualistic (left) and antagonistic (right) communities. As an example, we show restoration starting at 80% or 65% habitat loss for mutualistic and antagonistic communities, respectively. Top: fraction of interactions (across all mutualistic or antagonistic networks) for which either the ‘random’ (blue) or the ‘nonrandom’ (orange) strategy produces higher regional abundance, or there is no substantial difference between the two strategies (grey), at a given fraction of habitat loss. We assume that there is no difference between the two strategies when the abundance difference is less than 0.01. Middle: regional abundance of one interaction during habitat destruction (grey) and restoration (blue and orange). Bottom: example landscapes during restoration simulations shown in the middle panels when habitat loss is either 45% or 55%. Destroyed patches are coloured grey, patches where the interaction is present are black, and pristine but unoccupied patches are shown in white. Landscapes with blue and orange borders correspond to ‘random’ and ‘nonrandom’ restoration strategies, respectively.

## 4 Discussion

Adopting the metacommunity approach enabled us to explore the dynamics of community recovery during habitat restoration. Specifically, we investigated how the response of communities is affected by: (1) the amount of previously destroyed habitat, (2) the structure of the metacommunity, and (3) the spatial restoration strategy.

First, we found that the response of communities to restoration depends on how damaged the landscape is. When restoring relatively intact landscapes with large, connected areas of pristine habitats and evenly distributed areas of destroyed habitat, communities tend to recover their interaction richness and composition, and the interactions between species tend to recover their abundance across the landscape. In this case, the same outcome can be achieved irrespective of the order in which habitat patches are restored. This is because restoring any patch has the effect of increasing the available habitat area, rather than drastically improving the connectivity across the landscape.

However, when restoring highly fragmented landscapes, where pristine areas are small and disconnected, and communities may be at the brink of extinction, community recovery tends to be more difficult. Meta-analyses of forest restoration outcomes have also found less success in landscapes which were more fragmented prior to restoration (Crouzeilles et al., 2016, 2019). We found a lag in the recovery of the local interaction richness. This means that, the local networks are a smaller subset of the metanetwork during restoration than at the equivalent fraction of habitat loss during destruction. This is partly due to species becoming extinct, and partly to the spatial arrangement of pristine patches in the fragmented landscape. In the latter case, many pristine sites may be initially inaccessible, and thus, restoring more habitat creates corridors and allows these sites to be recolonised (Tilman et al., 1997; Gawecka and Bascompte, 2021). Differences in the structure and complexity of networks between restored and undisturbed sites have also been observed in restoration projects (Ribeiro da Silva et al., 2015; O’Connell et al., 2022).

In terms of the recovery of the regional abundance of interactions, we found contrasting responses of mutualistic and antagonistic communities when restoring highly destroyed landscapes. The former exhibit a lag, meaning that the regional abundance during restoration is lower than during destruction. This is again due to the fragmented arrangement of the pristine habitats. Conversely, antagonistic interactions experience a boost where their regional abundance is higher during restoration than destruction. When habitat is destroyed, the resources are released from consumer control due to the consumers becoming less abundant or even extinct (Kruess and Tscharntke, 1994; Liao et al., 2017). This reduced pressure allows them to achieve higher abundances during restoration. The increased abundance of resources benefits the remaining consumers, and thus, the abundance of their interactions is higher. However, it should be noted that this boost in abundance comes at the cost of loosing consumer species and interaction richness, with possible detrimental consequences on ecosystem functioning (Ritchie and Johnson, 2009; Schmitz et al., 2010; Hammerschlag et al., 2019).

Second, we found that the magnitude of the restoration lag or boost depends on the properties of the mentanetwork or the interaction. In mutualism, networks which are larger, more connected and more nested tend to have smaller restoration lags than the smaller and more modular networks. This means that identifying the topology of the metanetwork of mutualistic interactions may give us an indication of how the metacommunity will recover, and suggests that more care should be taken when restoring small or highly modular communities. In the case of antagonistic interactions, we found that the boost increases with the increasing interaction degree. In other words, interactions between two generalist species benefit more from restoration than highly specialised interactions, which indicates that more attention should be given to the latter when designing and monitoring restoration projects. This result echoes previous findings of specialist species being more sensitive to habitat loss and fragmentation than generalists (Aizen et al., 2012; Emer et al., 2018; Li et al., 2022).

Lastly, restoration efficiency depends on the order in which habitat patches are restored. Our results show that prioritising restoration of patches adjacent to the most species-rich communities (‘nonrandom’ restoration strategy) increases the overall efficiency of restoration, especially in the case of antagonistic communities. This is consistent with previous findings on the response of a single species and simple communities with two to four species (Huxel and Hastings, 1999; Gawecka and Bascompte, 2021). However, in mutualism, we found that the efficiency may be increased further by adopting a two-stage restoration strategy: initially restoring patches adjacent to occupied ones, followed by restoring patches at random. When the landscape is highly fragmented and very few patches harbour communities, restoration should be carried out in the proximity to these communities, creating a large cluster of pristine sites. This may bring the species back from the brink of extinction and allow them to achieve viable population sizes. However, continuing restoration in this manner may at some point prevent recolonisation of other parts of the landscape. Restoring patches in a random order instead creates numerous small clusters of pristine sites, and thus, improves the overall connectivity of the landscape. This is in agreement with a recent study which, by synthesising theoretical concepts and empirical evidence, recommend that forest cover should be configured in one large patch and many evenly distributed small patches (Arroyo-Rodríguez et al., 2020). The significance of continuous habitats connected by corridors has been highlighted in a number of studies (Newmark et al., 2017; Thompson et al., 2017; Hyseni et al., 2021; San-José et al., 2022).

The importance of the spatial structure of the landscape depends on the species’ dispersal abilities (King and With, 2002; Grilli et al., 2015). As our model assumes dispersal to the nearest neighbouring patches, our findings are particularly applicable to communities composed of species which tend to disperse locally. Species which are able to traverse long distances may be affected by fragmentation to a lesser extent, and thus we expect smaller restoration lags and differences between spatial restoration strategies (Forup et al., 2008; Emer et al., 2018; Gawecka and Bascompte, 2021). Furthermore, although our model considers a binary description of patch state (pristine or destroyed), both habitat destruction and restoration may involve gradual changes in habitat quality (Banks-Leite et al., 2020). Given that habitat quality has an effect on the persistence and structure of ecological communities (Moilanen and Hanski, 1998; Thomas et al., 2001; Chisholm et al., 2011; Miller and Allesina, 2021; Silva et al., 2022), it is likely to affect their response to restoration.

The current state of our ecosystems is dismal. For example, 70% of world’s forest are located within one kilometre of the forest’s edge (Haddad et al., 2015), and the majority of forests are at the tipping point between continuous and fragmented states (Taubert et al., 2018). Thus, restoration of our ecosystems is now more urgent than ever. We showed that the recovery of communities in fragmented landscapes is dependent on the structure of these species interaction networks. Moreover, we demonstrated that the recovery may be enhanced by adopting certain spatial restoration strategies. Overall, our study highlights the importance of careful designing of restoration projects in fragmented landscapes.

## Acknowledgements

We thank the members of Bascompte Lab for discussions. Funding was provided by University of Zurich Postdoc Grant (grant number FK-22-114 to KAG) and the SNSF (grant number 310030_197201 to JB).

## Data availability

All code used in this study is available on GitHub (github.com/kgawecka/habitat_restoration_networks).

