## Supporting Information for "Habitat restoration and the recovery of metacommunities"

Table S1: Empirical networks used in this study (M - mutualistic, A - antagonistic, PL - plant-pollinator, SD - plant-seed disperser, HP - host-parasite, PH - plant-herbivore; available at www.web-of-life.es.)

| **network** | **resources** | **consumers** | **interactions** | **reference** |
|---|---|---|---|---|
| M_PL_006 | 17 | 61 | 146 | Dicks et al. (2002) |
| M_PL_010 | 31 | 76 | 456 | Elberling & Olesen (unpublished) |
| M_PL_025 | 13 | 44 | 143 | Motten (1986) |
| M_PL_033 | 13 | 34 | 141 | Small (1976) |
| M_PL_036 | 10 | 12 | 30 | Olesen (unpublished) |
| M_PL_037 | 10 | 40 | 72 | Montero (2005) |
| M_PL_039 | 17 | 51 | 129 | Stald (2003) |
| M_PL_046 | 16 | 44 | 278 | Bundgaard (2003) |
| M_PL_051 | 14 | 90 | 104 | Vázquez (2003) |
| M_PL_059 | 13 | 13 | 71 | Bezerra et al. (2009) |
| M_SD_002 | 31 | 9 | 119 | Beehler (1983) |
| M_SD_005 | 25 | 13 | 49 | Carlo et al. (2003) |
| M_SD_007 | 72 | 7 | 143 | Crome (1975) |
| M_SD_008 | 16 | 10 | 110 | Frost (1980) |
| M_SD_010 | 50 | 14 | 234 | Snow and Snow (1971) |
| M_SD_012 | 35 | 29 | 146 | Galetti and Pizo (1996) |
| M_SD_014 | 16 | 17 | 121 | Jordano (1985) |
| M_SD_016 | 24 | 61 | 500 | Lambert (1989) |
| M_SD_025 | 7 | 6 | 22 | Sorensen (1981) |
| M_SD_027 | 12 | 4 | 31 | Jordano (unpublished) |
| A_HP_002 | 18 | 24 | 96 | Hadfield et al. (2014) |
| A_HP_005 | 7 | 13 | 51 | Hadfield et al. (2014) |
| A_HP_007 | 8 | 17 | 43 | Hadfield et al. (2014) |
| A_HP_008 | 8 | 24 | 37 | Hadfield et al. (2014) |
| A_HP_012 | 7 | 23 | 63 | Hadfield et al. (2014) |
| A_HP_015 | 3 | 7 | 12 | Hadfield et al. (2014) |
| A_HP_021 | 9 | 15 | 57 | Hadfield et al. (2014) |
| A_HP_025 | 18 | 40 | 107 | Hadfield et al. (2014) |
| A_HP_028 | 4 | 15 | 19 | Hadfield et al. (2014) |
| A_HP_032 | 14 | 13 | 32 | Hadfield et al. (2014) |
| A_HP_033 | 22 | 25 | 198 | Hadfield et al. (2014) |
| A_HP_035 | 6 | 7 | 15 | Hadfield et al. (2014) |
| A_HP_042 | 21 | 32 | 84 | Hadfield et al. (2014) |
| A_HP_046 | 17 | 39 | 202 | Hadfield et al. (2014) |
| A_HP_047 | 11 | 26 | 100 | Hadfield et al. (2014) |
| A_HP_050 | 27 | 35 | 226 | Hadfield et al. (2014) |
| A_PH_004 | 52 | 22 | 184 | Joern (1979) |
| A_PH_005 | 54 | 24 | 173 | Joern (1979) |
| A_PH_006 | 6 | 88 | 116 | Leather (1991) |
| A_PH_007 | 5 | 64 | 95 | Leather (1991) |

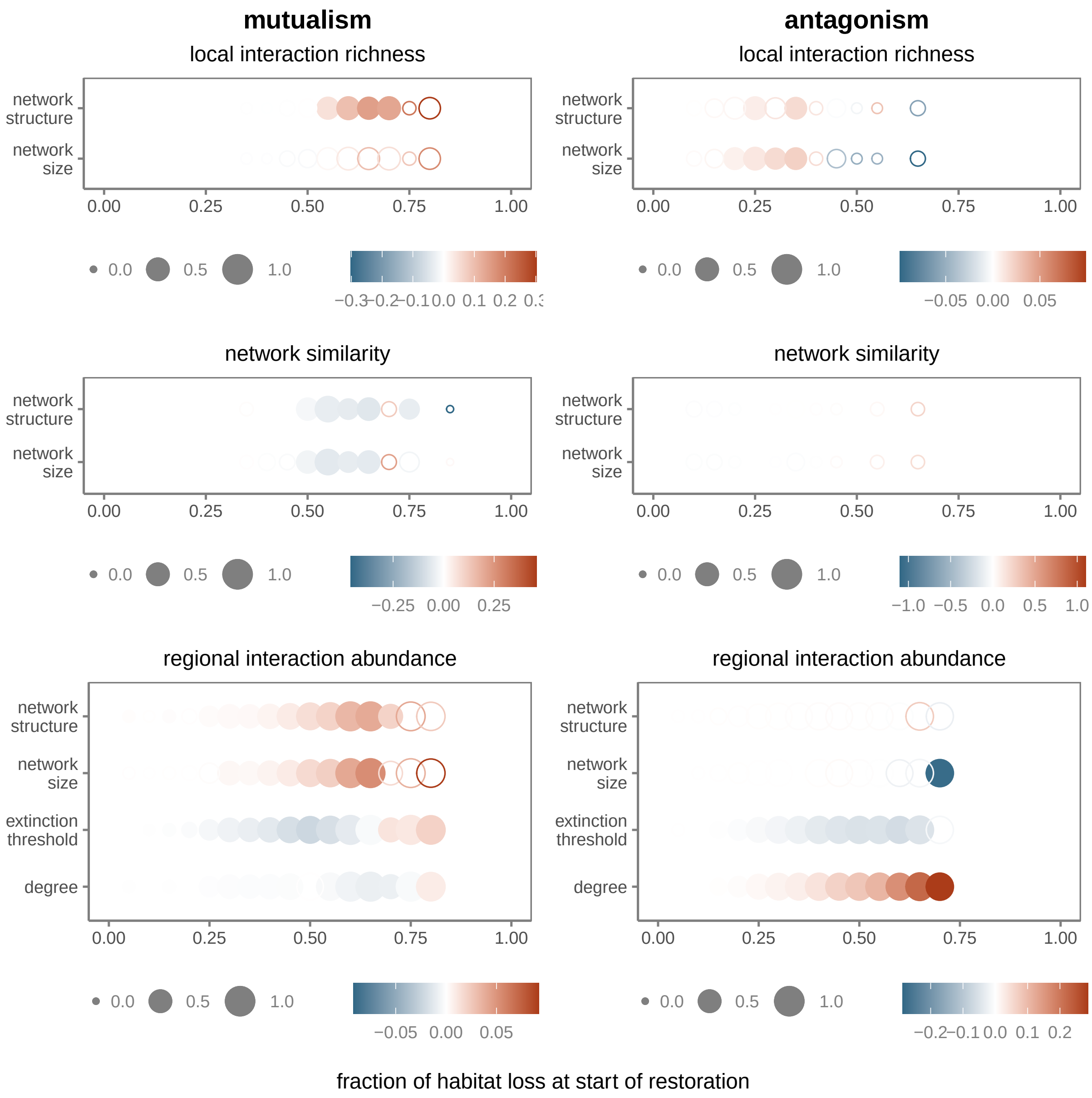


Figure S1: The effect of metanetwork and interaction properties on restoration efficiency of mean local interaction richness (top), mean similarity of local networks (middle), and regional interaction abundance (bottom). For local interaction richness and network similarity, we fitted linear models with metanetwork structure and metanetwork size as the explanatory variables. For regional interaction abundance, we fitted linear mixed models with metanetwork structure, metanetwork size, interaction extinction threshold and interaction degree as the explanatory variables. We fitted the models to the results of simulations with mutualistic (left) and antagonistic (right) networks, and 'nonrandom' restoration starting at different fractions of habitat loss. The size of the points indicates the $R^2$, full and empty circles correspond to significant (p-value $< 0.01$) and non-significant effects, respectively, and the colours show the slope parameter estimate. The explanatory variables are scaled, meaning that the magnitudes of their slope parameter estimates are comparable for each interaction type and each restoration simulation.

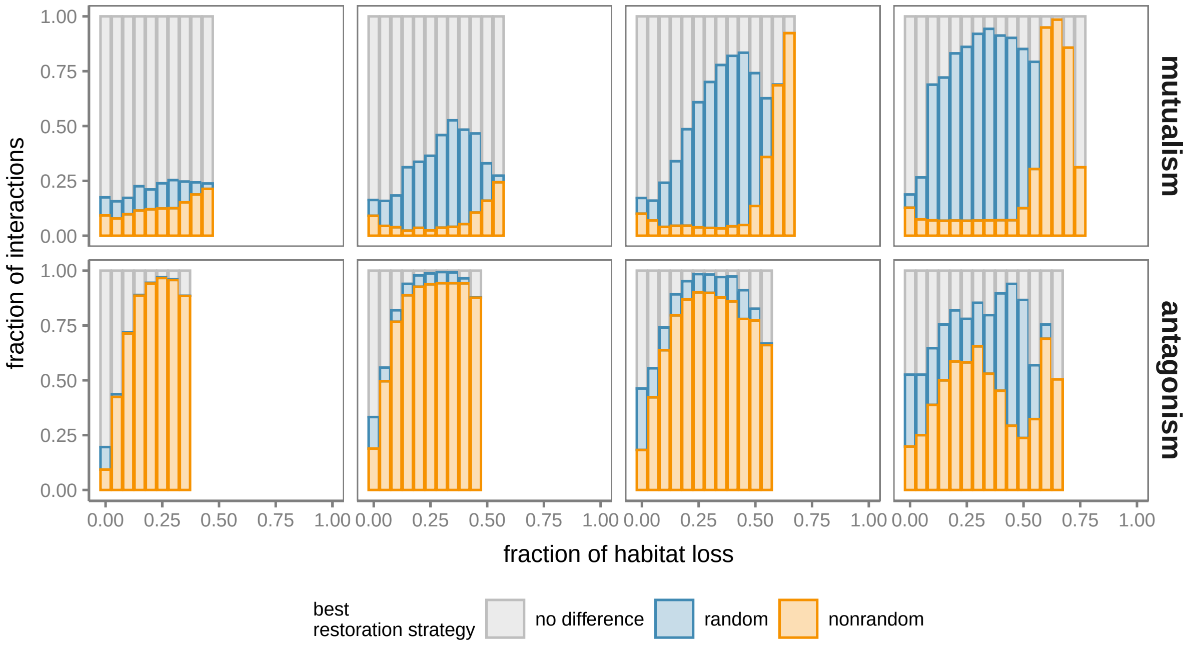


Figure S2: Fraction of interactions (across all mutualistic or antagonistic networks) for which either the 'random' (blue) or the 'nonrandom' (orange) strategy produces higher regional abundance, or there is no substantial difference between the two strategies (grey), at a given fraction of habitat loss. We assume that there is no difference between the two strategies when the abundance difference is less than 0.01. Panels show restoration simulations starting at different fractions of habitat loss (from left to right: 50, 60, 70 and 80% for mutualistic networks, and 40, 50, 60 and 70% for antagonistic networks).

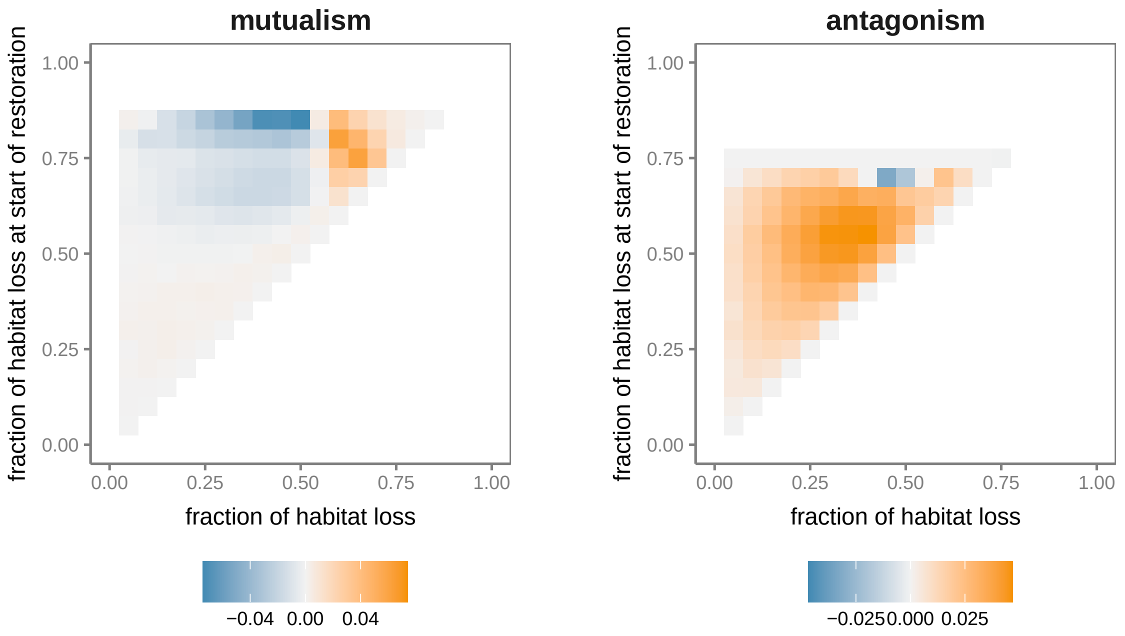


Figure S3: Differences in regional interaction abundance between ‘random’ and ‘nonrandom’ restoration strategies at different fractions of habitat loss (horizontal axis) in restoration simulations starting at different fractions of habitat loss (vertical axis). Grid cell colour shows the median difference across all interactions in either mutualistic (left) or antagonistic (right) networks. Positive difference values (orange) indicate that the ‘nonrandom’ strategy produced higher abundance. Negative difference values (blue) indicate that the ‘random’ strategy produced higher abundance.

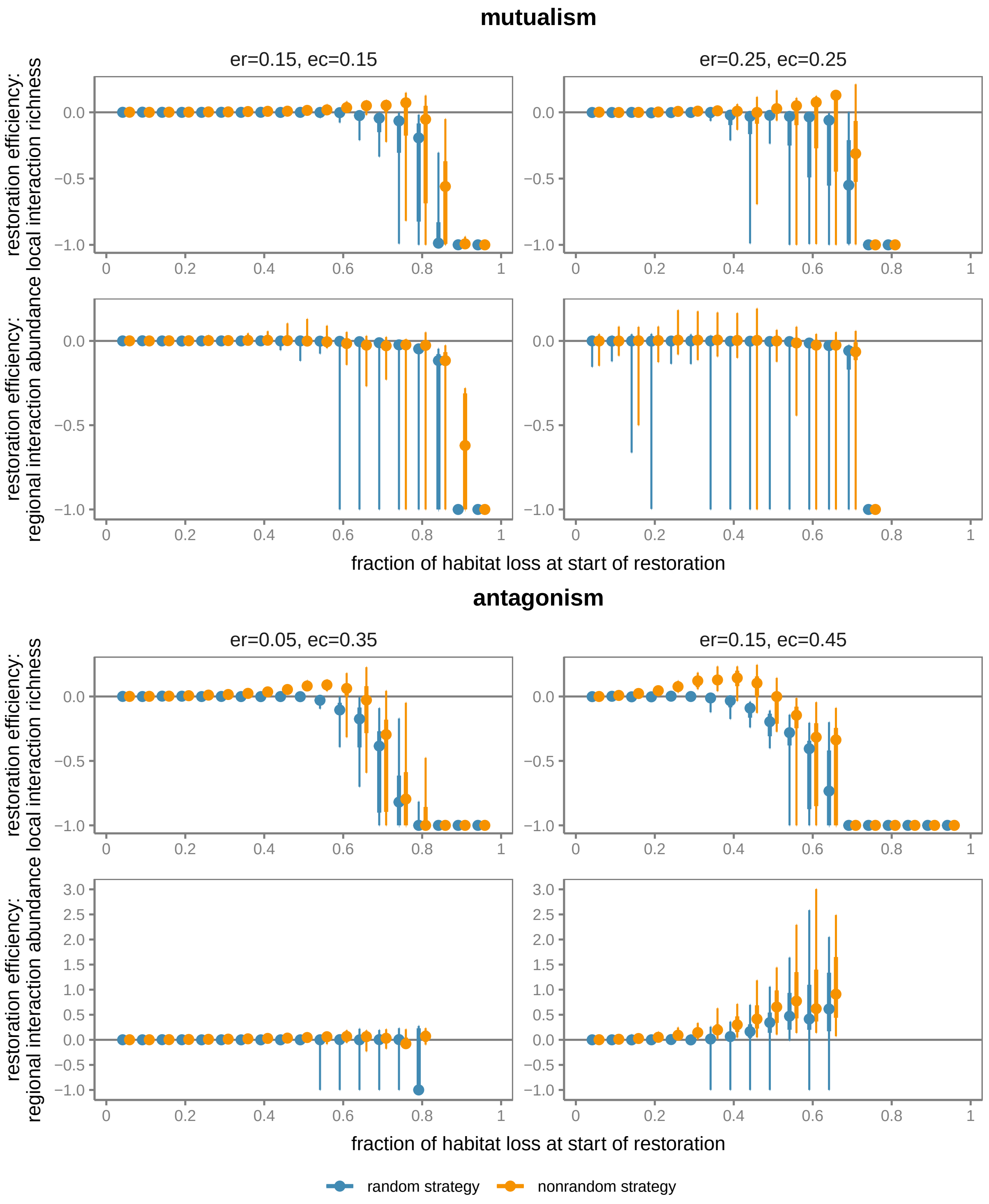


Figure S4: Effect of model parameters on restoration efficiency trends. Plots show restoration efficiency of mean local interaction richness and regional interaction abundance as a function of the fraction of habitat loss at which restoration begins. Left and right panels correspond to simulations with lower and higher intrinsic extinction probabilities, respectively, than those presented in the main text. Points, thicker vertical lines and thinner vertical lines represent the median, interquartile range, and minimum and maximum values, respectively, across all mutualistic or antagonistic networks and interactions. Random and nonrandom restoration strategies are shown in blue and orange, respectively. Positive restoration efficiency values indicate a ‘restoration boost’ (i.e., the quantity is, on average, higher during restoration than during destruction), whereas negative restoration efficiency values imply a ‘restoration lag’ (i.e., the quantity is, on average, lower during restoration than during destruction).

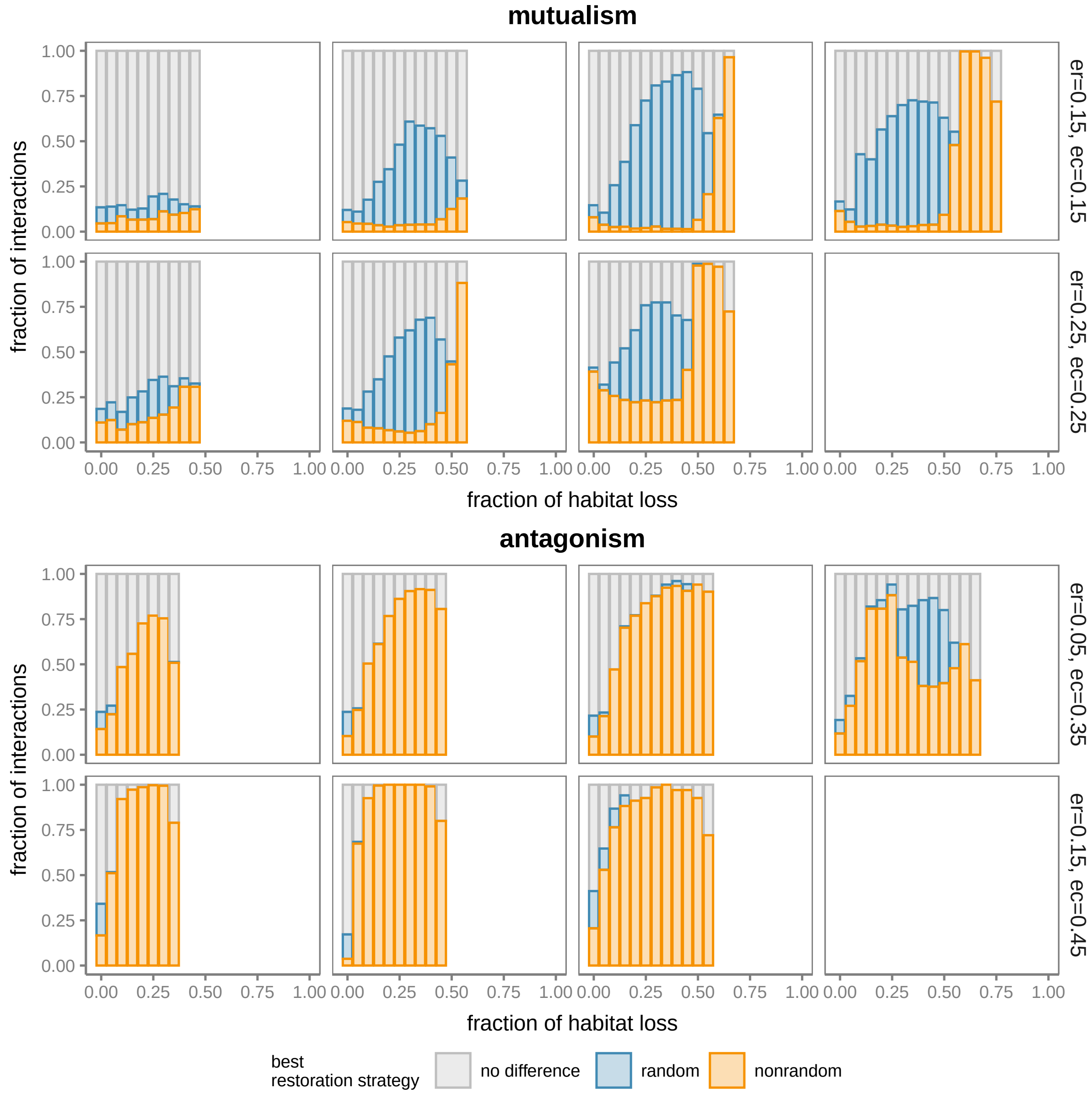


Figure S5: Effect of model parameters on the most efficient restoration strategy. Plots show fraction of interactions (across all mutualistic or antagonistic networks) for which either the 'random' (blue) or the 'nonrandom' (orange) strategy produces higher abundance, or there is no substantial difference between the two strategies (grey), at a given fraction of habitat loss. We assume that there is no difference between the two strategies when the abundance difference is less than 0.01. Upper and lower panels correspond to simulations with lower and higher intrinsic extinction probabilities, respectively, than those presented in the main text. Panels show restoration simulations starting at different fractions of habitat loss (from left to right: 50, 60, 70 and 80% for mutualistic networks, and 40, 50, 60 and 70% for antagonistic networks).

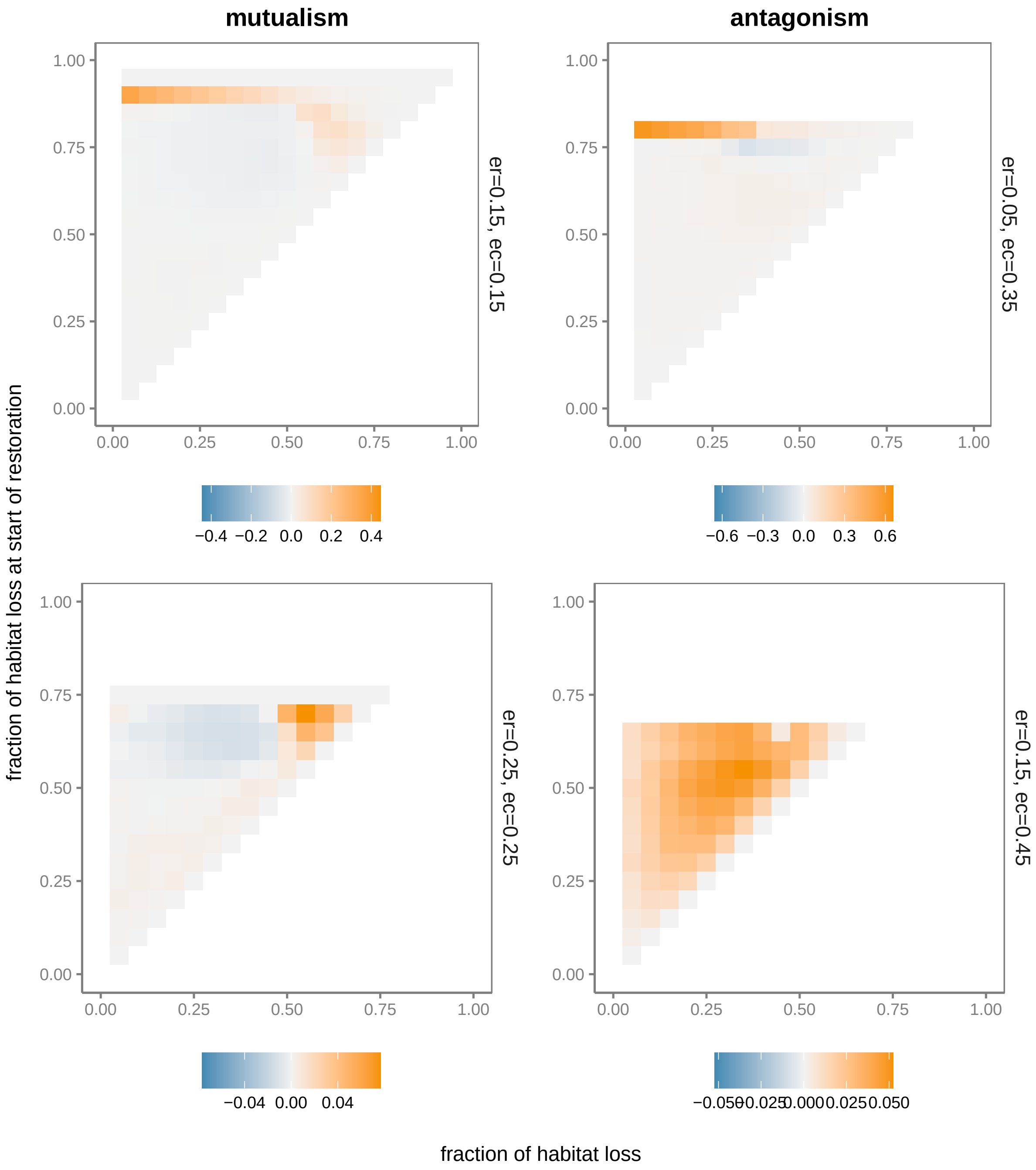


Figure S6: Effect of model parameters on differences in regional interaction abundance between 'random' and 'nonrandom' restoration strategies. Plots show different fractions of habitat loss (horizontal axis) in restoration simulations starting at different fractions of habitat loss (vertical axis). Grid cell colour shows the median difference across all interactions in either mutualistic (left) or antagonistic (right) networks. Positive difference values (orange) indicate that the 'nonrandom' strategy produced higher abundance. Negative difference values (blue) indicate that the 'random' strategy produced higher abundance. Upper and lower panels correspond to simulations with lower and higher intrinsic extinction probabilities, respectively, than those presented in the main text.

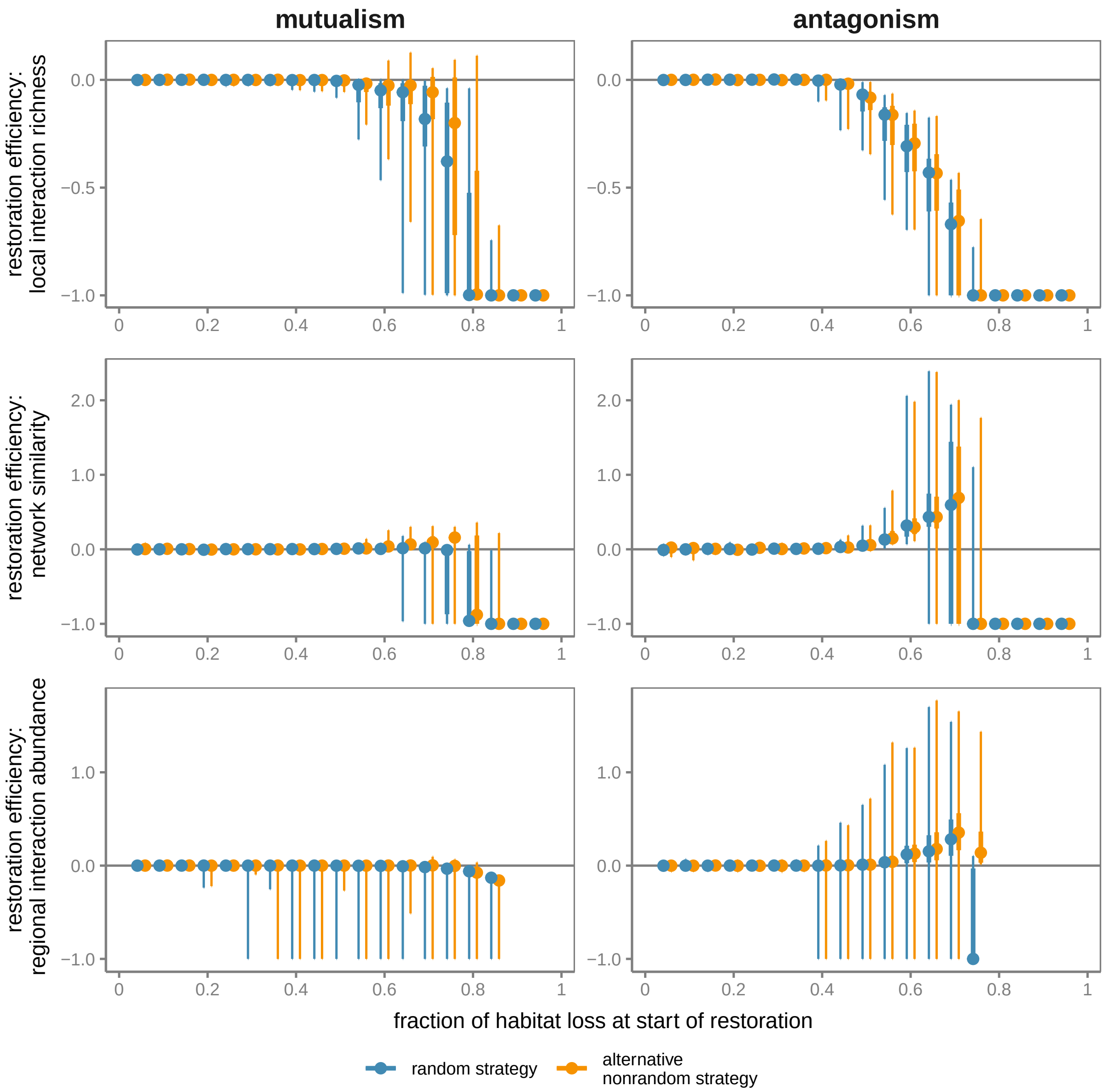


Figure S7: Restoration efficiency as a function of the fraction of habitat loss at which restoration begins. The panels show restoration efficiency of: mean local interaction richness (top), mean similarity of local networks (middle), and regional interaction abundance (bottom). Points, thicker vertical lines and thinner vertical lines represent the median, interquartile range, and minimum and maximum values, respectively, across all mutualistic (left) or antagonistic (right) networks and interactions. 'random' and 'alternative nonrandom' restoration strategies are shown in blue and orange, respectively. Positive restoration efficiency values indicate a 'restoration boost' (i.e., the quantity is, on average, higher during restoration than during destruction), whereas negative restoration efficiency values imply a 'restoration lag' (i.e., the quantity is, on average, lower during restoration than during destruction).

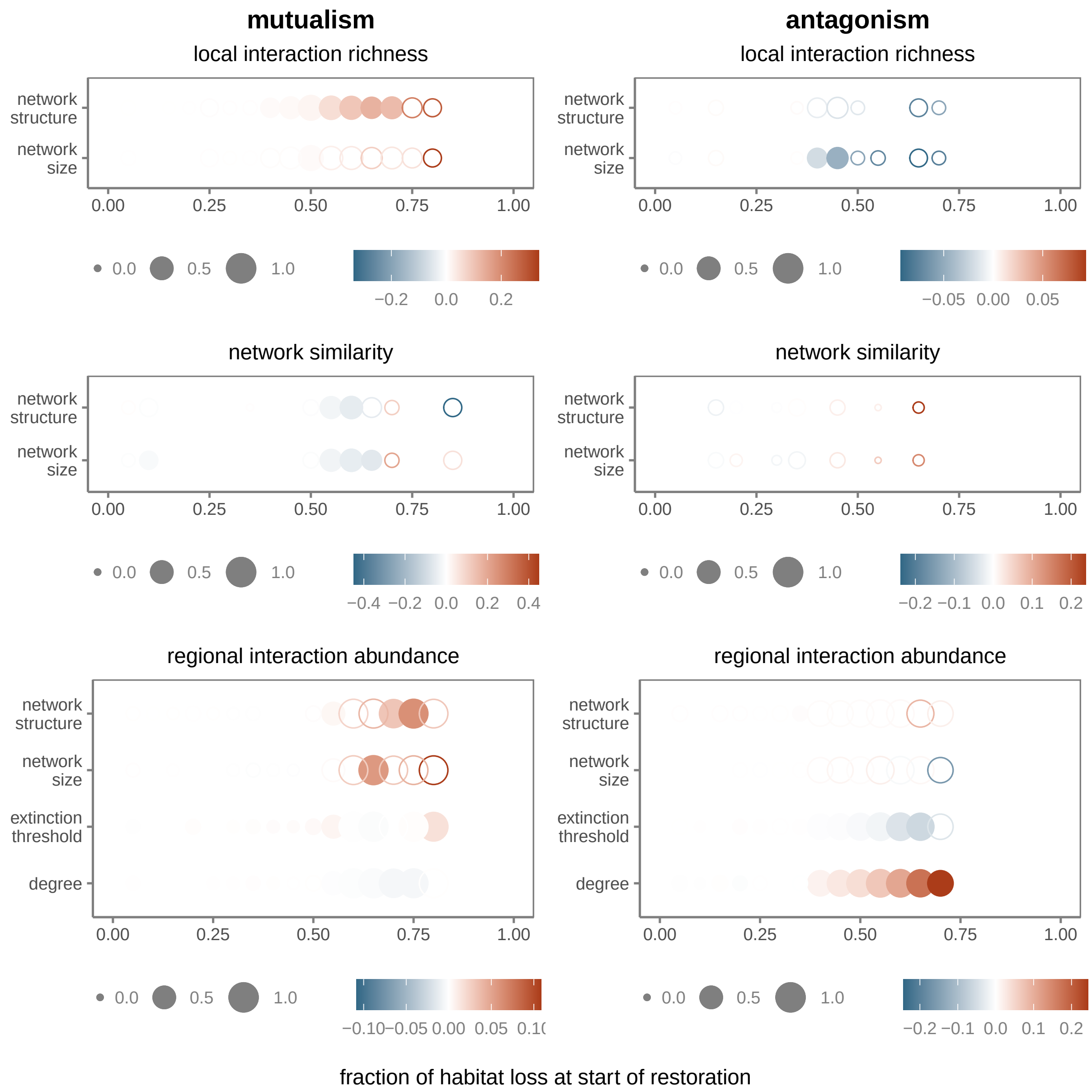


Figure S8: The effect of metanetwork and interaction properties on restoration efficiency of mean local interaction richness (top), mean similarity of local networks (middle), and regional interaction abundance (bottom). For local interaction richness and network similarity, we fitted linear models with metanetwork structure and metanetwork size as the explanatory variables. For regional interaction abundance, we fitted linear mixed models with metanetwork structure, metanetwork size, interaction extinction threshold and interaction degree as the explanatory variables. We fitted the models to the results of simulations with mutualistic (left) and antagonistic (right) networks, and 'alternative nonrandom' restoration starting at different fractions of habitat loss. The size of the points indicates the $R^2$, full and empty circles correspond to significant (p-value $<$ 0.01) and non-significant effects, respectively, and the colours show the slope parameter estimate. The explanatory variables are scaled, meaning that the magnitudes of their slope parameter estimates are comparable for each interaction type and each restoration simulation.

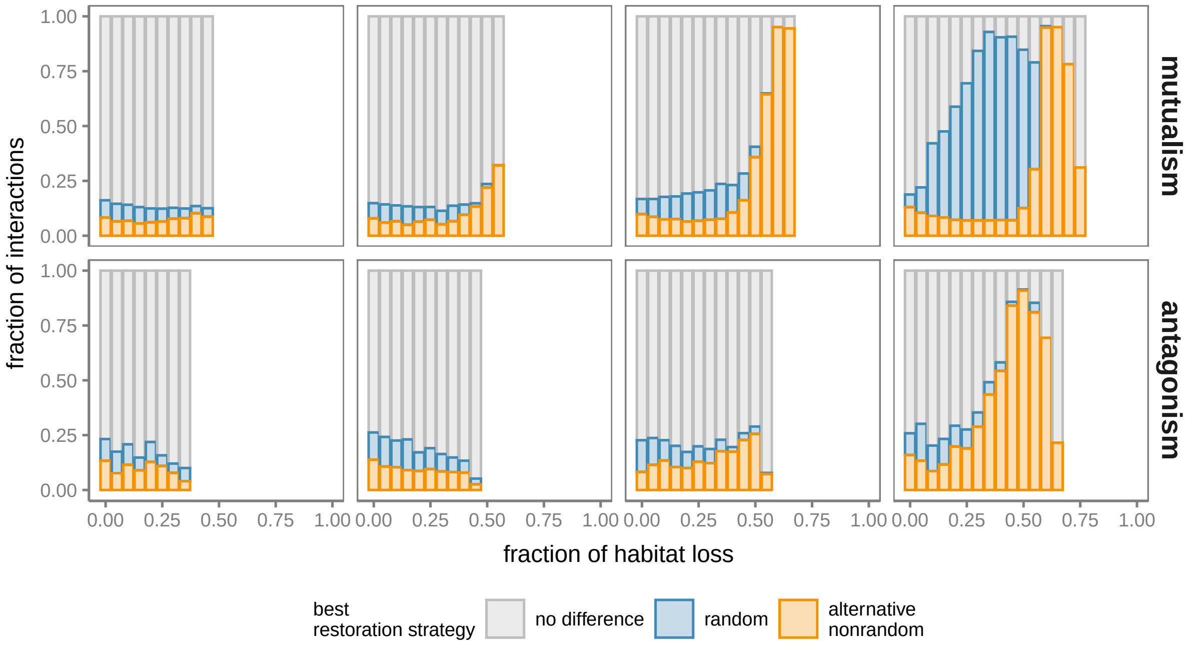


Figure S9: Fraction of interactions (across all mutualistic or antagonistic networks) for which either the 'random' (blue) or the 'alternative nonrandom' (orange) strategy produces higher abundance, or there is no substantial difference between the two strategies (grey), at a given fraction of habitat loss. We assume that there is no difference between the two strategies when the abundance difference is less than 0.01. Panels show restoration simulations starting at different fractions of habitat loss (from left to right: 50, 60, 70 and 80% for mutualistic networks, and 40, 50, 60 and 70% for antagonistic networks).

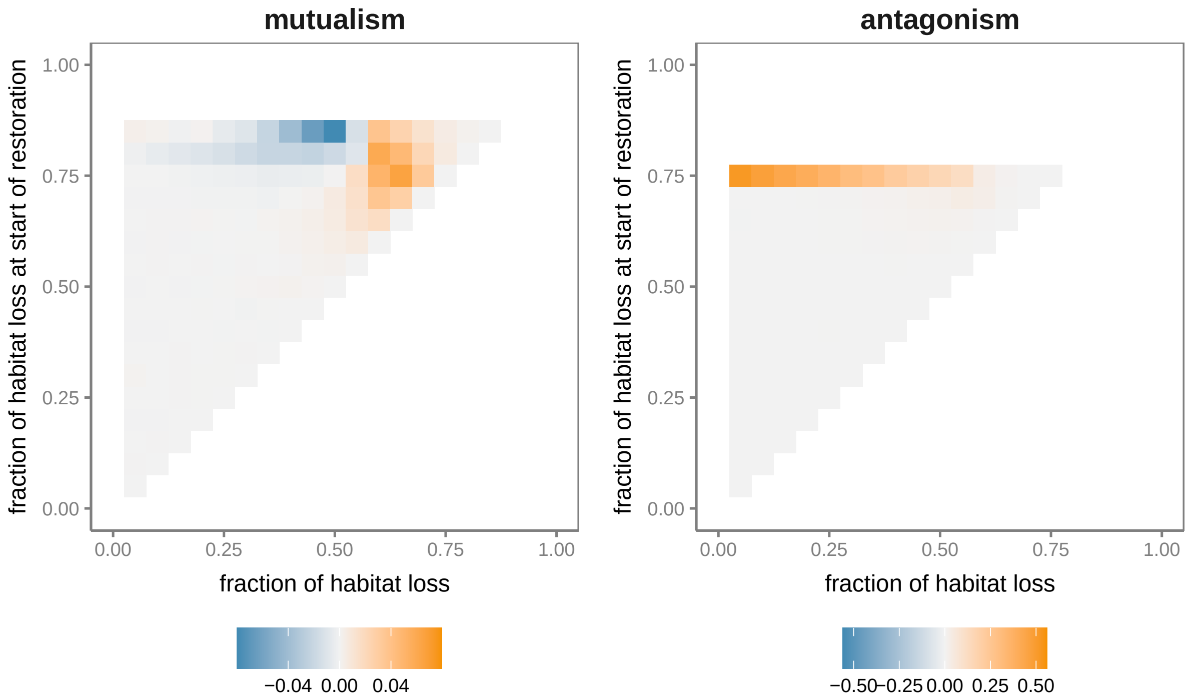


Figure S10: Differences in regional interaction abundance between 'random' and 'alternative nonrandom' restoration strategies at different fractions of habitat loss (horizontal axis) in restoration simulations starting at different fractions of habitat loss (vertical axis). Grid cell colour shows the median difference across all interactions in either mutualistic (left) or antagonistic (right) networks. Positive difference values (orange) indicate that the 'alternative nonrandom' strategy produced higher abundance. Negative difference values (blue) indicate that the 'random' strategy produced higher abundance.
